# Technical Note: Quantifying music-dance synchrony with the application of a deep learning-based 2D pose estimator

**DOI:** 10.1101/2020.10.09.333617

**Authors:** Filip Potempski, Andrea Sabo, Kara K Patterson

## Abstract

Dance interventions are more effective at improving gait and balance outcomes than other rehabilitation interventions. Repeated training may culminate in superior motor performance compared to other interventions without synchronization. This technical note will describe a novel method using a deep learning-based 2D pose estimator: OpenPose, alongside beat analysis of music to quantify movement-music synchrony during salsa dancing. This method has four components: i) camera setup and recording, ii) tempo/downbeat analysis and waveform cleanup, iii) OpenPose estimation and data extraction, and iv) synchronization analysis. Two trials were recorded: one in which the dancer danced synchronously to the music and one where they did not. The salsa dancer performed a solo basic salsa step continuously for 90 seconds to a salsa track while their movements and the music were recorded with a webcam. This data was then extracted from OpenPose and analyzed. The mean synchronization value for both feet was significantly lower in the synchronous condition than the asynchronous condition, indicating that this is an effective means to track and quantify a dancer’s movement and synchrony while performing a basic salsa step.

## Introduction

Dance is a universal human activity that confers many benefits including aerobic fitness, better balance, emotional and social well-being, and stress reduction.^1,2^ Anyone can benefit from dance including trained dancers, recreational dancers, older adults, and people with neurological conditions including Parkinson’s disease and stroke.^3–10^ As an adjunct to rehabilitation, dance interventions are more effective at improving gait and balance outcomes than other rehabilitation interventions.^11^

The advantages of dance over conventional exercise interventions may be derived from the synchronized movement to the rhythmic cues embedded in music. The extensive connectivity between the auditory and motor systems forms the basis for entrainment between rhythmic auditory signals and motor responses such as tapping your foot along to the beat in music.^12–18^ These spontaneous synchronized movements may result from the processing of perceived rhythms by motor areas in the brain including the basal ganglia, supplementary motor area.^16,19,20^ Thus, during dance, the rhythmic cues embedded in music, combined with the goal of synchronizing movement to those cues, may drive motor output. Repeated training may culminate in superior motor performance (such as better balance control) compared to other interventions without synchronization. This proposed mechanism is supported by the fact that music enhances motor performance and reduces metabolic costs of exercise in healthy adults^12,21–23^ and rhythmic auditory cueing during rehabilitation sessions improves gait performance^24^ and brain activation patterns^25^ post-stroke.

To investigate this potential mechanism, it will be necessary to measure the movement of people while they are dancing and quantify how well they synchronize their movement to the music. The accuracy of this type of measurement depends heavily on the device used as well as the location of that device on the body.^20,26^ Quantitative movement analysis often requires costly devices such as 3D capture systems, accelerometers, gyroscopes, or force plates^27,28^. However, vision-based tracking systems are less expensive and less cumbersome alternatives.^29^ With this in mind, our group used OpenPose,^30,31^ a deep learning-based 2D keypoint estimator, to track dancer movements and subsequently compare the timing of their footfall events to the timing of the beat of the music to quantify movement synchronization. OpenPose^30,31^ has been used for many different objectives including 3D pose estimation,^32^ 3D model generation,^33^ and analyzing features of gait.^34^ This open-source software automatically obtains joint coordinates of the individuals in the image or video, enabling the calculation of parameters of interest. In this technical note, we will describe our novel method using OpenPose ^30,31^ alongside beat analysis of music to quantify movement-music synchrony during salsa dancing.

## Methods

Our method quantifies how well a dancer’s steps synchronize with the downbeat of the music to which they are dancing. We used 1 webcam (Logitech Meetup^35^, 1920×1080 pixels, 30 fps) 1 desktop computer (Windows PC), and computer speakers in this process. A recreational salsa dancer (5 years experience) performed a solo basic salsa step continuously for 90 seconds to a salsa track while her movements and the salsa track (1411 Kbps) were recorded with the webcam. The method and analysis have four components: i) camera setup and video recording of salsa dancing, ii) tempo/downbeat analysis and waveform cleanup, iii) OpenPose estimation and data extraction, and iv) synchronization analysis. Figure 1 provides a visual summary of the data flow and interaction between these components.

**Figure 1.**
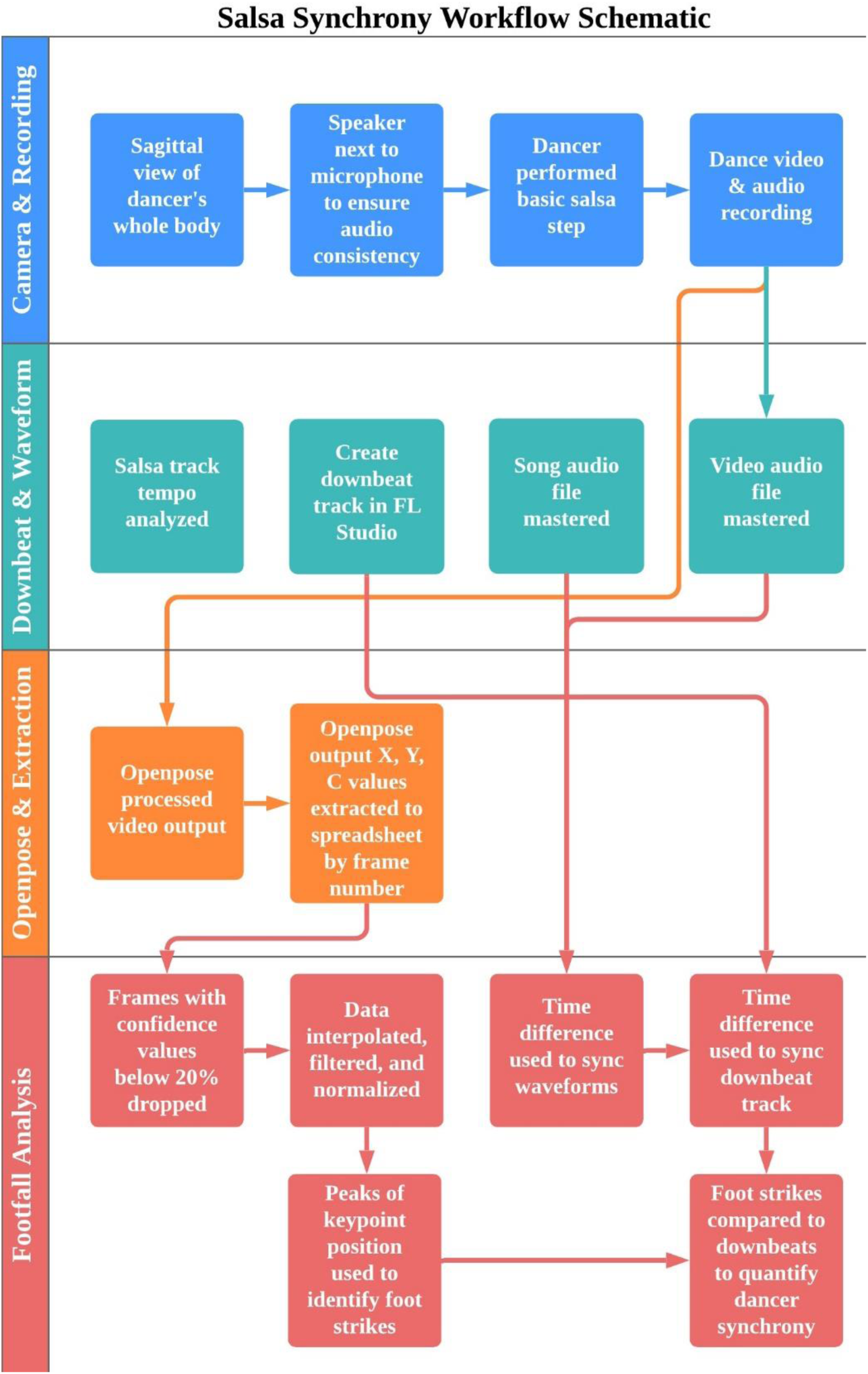
Workflow for salsa synchrony measure. Abbreviations: FL = Fruity loops; X = x coordinates, Y = y coordinates, C = confidence values.

### Camera Setup and Video Recording of Salsa Dancing

The webcam was positioned to capture the dancer’s whole body in a sagittal view and to avoided filming at an angle to reduce image distortion (dance video). This is important to avoid parallax error in lateral views.^36^ We also reduced visual background clutter and ensured the participant was well lit to avoid further information inaccuracies during OpenPose analysis. We placed the speaker playing the salsa track next to the microphone of the webcam to ensure consistency between audio waveforms of the salsa track and the dance video. We recorded audio on the same device as the video. Our participant performed a basic salsa step (Figure 2), continuously for 90 seconds under two conditions. In the synchronous condition, she danced on in time with the music. In the asynchronous condition, she intentionally danced out of time with the music.

**Figure 2.**
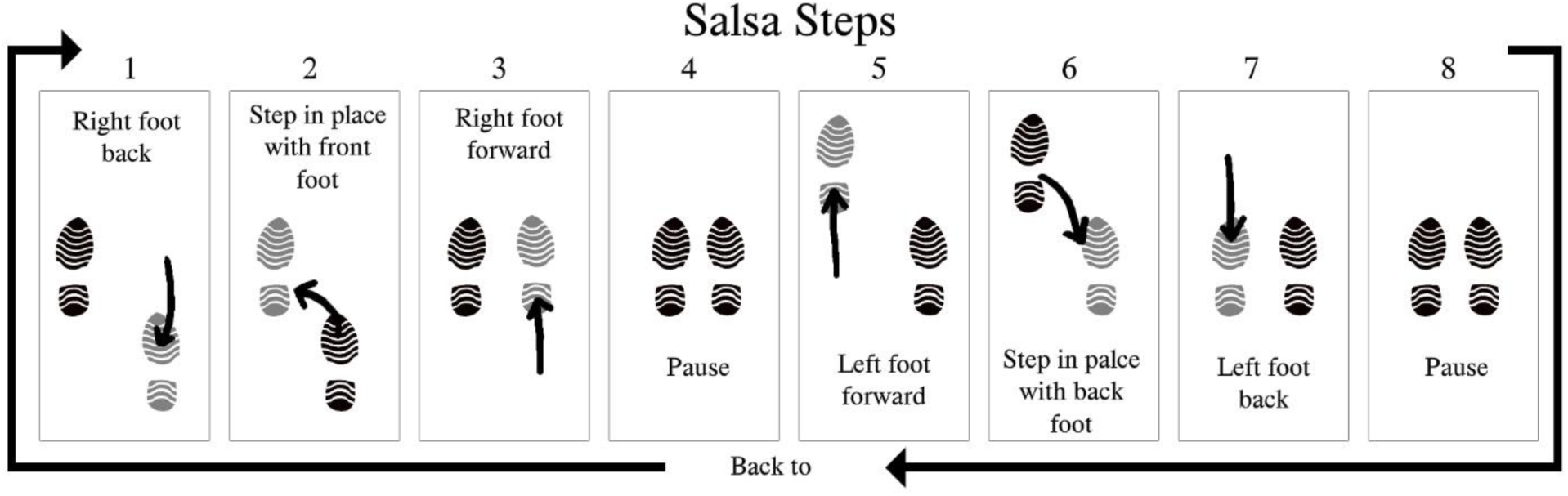
Salsa Step Pattern. The positioning (represented by shoe prints) and direction (represented by arrows) of steps for the basic salsa step. The music counts on which each step occurs are listed above.

### Tempo/Downbeat Analysis and Waveform Cleanup

Separate from the video recording of the salsa dancing, we analyzed the tempo of the salsa track with the free version of Fruity Loops (FL) Studio (Edition 20.6, Image Line Software, Ghent, Belgium) digital audio workstation (DAW) and its built-in tempo analysis. From this analysis, we created a new audio track of audible beats matched to the salsa song (downbeat track). We used a tone for the audible beats with limited distortion to ensure a clean audio waveform for later analysis. We visually and aurally evaluated the downbeat track to ensure it aligned with the salsa track and exported the track as a .WAV file.

To clean up the audio waveforms recorded in the dance video as well as the song track for later analysis, we used FL Studio. We used the equalizer tool to cut all frequencies below 40Hz in both recordings to reduce background noise. We then used the multiband compressor tool to highlight high and low frequencies in the recordings to make the peaks and troughs of the waveforms more differentiable. We used a second equalizer tool to further boost relevant frequencies, followed by a limiter to prevent clipping the audio on the high end and distorting the waveform. Finally, we exported the mastered audio files as .WAV files for analysis.

### OpenPose Estimation and Data Extraction

A public release of OpenPose (Version 1.5.0) was run on a Windows machine and used to analyze the dance video. Each frame of the dance video was processed to output a .JSON file containing the X and Y coordinates and confidence values for each of the 25 estimated keypoints (Figure 2). These files were then zipped into a single folder. To avoid working with many small files, the data from the .JSON files were extracted to a .CSV file using a Python script and ordered as a function of frame number.

### Footfall & Salsa Synchrony Analysis

The .CSV file created in Python was imported to MATLAB (Version 9.7, Mathworks, Natick, MA, USA) where the keypoints defined by OpenPose, and corresponding to the big toe of the left and right feet (keypoints 19 and 22 respectively^37^) were used to create a new table. All frames with confidence values under 20% were dropped and data was interpolated using MATLAB’s *interp1*^38^ function, then NaN values were interpolated using the *inpaint_NaNs* function^39^ with a plate equation and an interpolating operator of delta cubed. Data was then filtered with a lowpass filter (2^nd^ order Butterworth with a cut-off frequency at 4 Hz) using the zero-phase filtering operation: *filtfilt*^40^ and normalized to −1,1 coordinates. The time of each frame was derived from the frame number over the recording device’s frames per second rate.

The audio waveforms from the dance video, salsa track, and downbeat track were also imported into MATLAB and lowpass filtered (2^nd^ order Butterworth with a cut-off frequency of 0.707 Hz). Then, using MATLAB’s *alignsignals*^41^ function, the salsa track waveform was synced up to the video waveform. Based on the time difference between the two waveforms, the downbeat track waveform was aligned to match up to the video waveform. Utilizing MATLAB’s *findpeaks*^42^ function, each peak of the downbeat track waveform was marked to represent the music’s downbeats.

Since the participant was recorded in a sagittal view most of the movement or interest was captured by the x-values. The peaks or troughs of the keypoint of interest’s X-value position were used to indicate foot strikes. Using MATLAB’s *findpeaks*^42^ function again, these peaks were found. We then overlaid the downbeat points onto the foot position data for a visual representation of movement synchrony as seen in Figure 3. Time, foot strike, and beats were combined into a single table and exported as an .XLS file for further analysis. Synchronization of the participant’s steps to the music was quantified by calculating the difference in time (seconds) between the occurrence of beats 1 and 5 of the music to the occurrence of the nearest right backward and left forward footfall events respectively (as described in Figure 2). These values were averaged for each foot over each of the two conditions (synchronous and asynchronous).

**Figure 3a/b.**
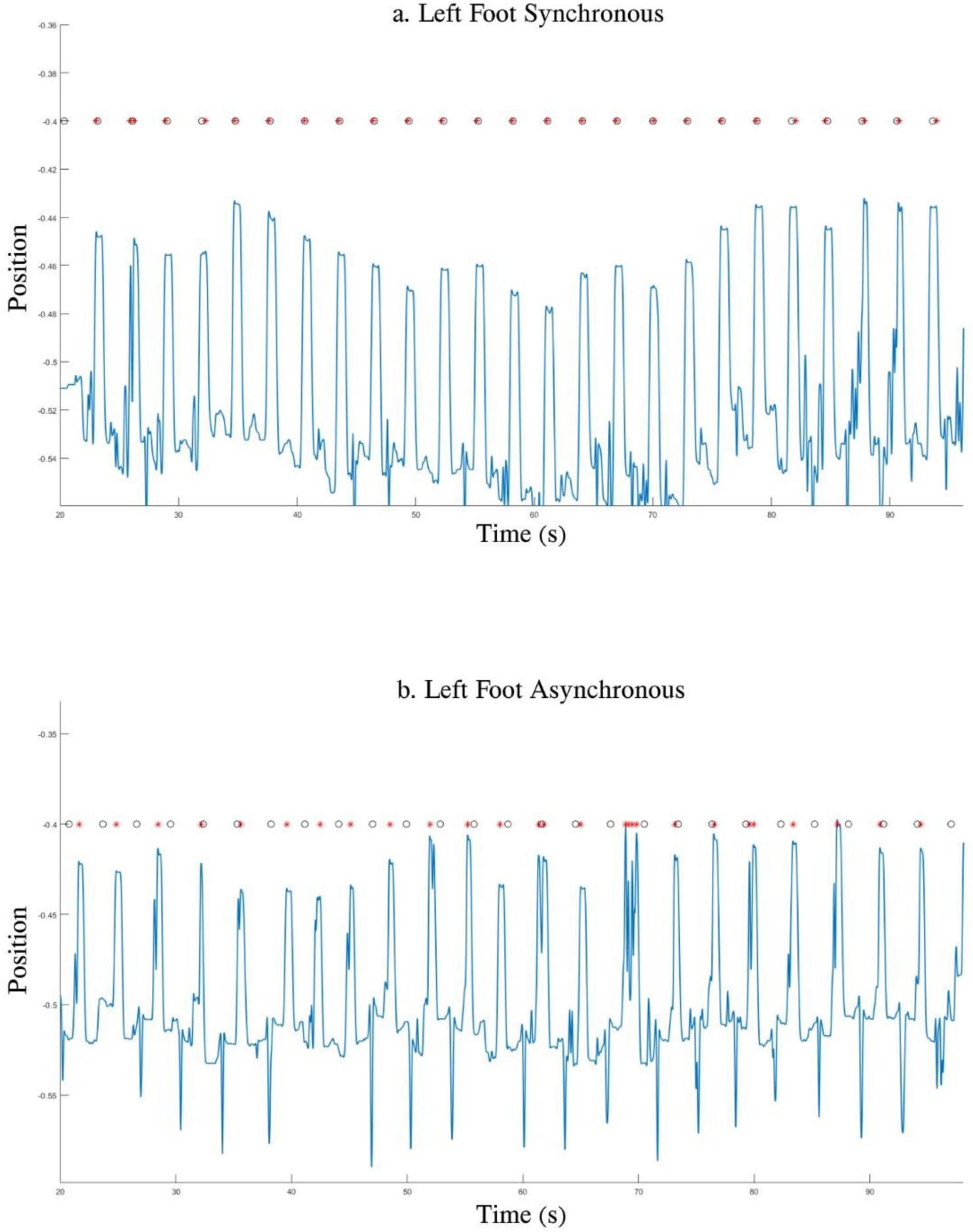
Plot of Estimated Synchronized & Asynchronized Left Foot Strikes against Downbeat 5. Black circles indicate downbeats, red stars indicate foot strikes, and blue lines indicate keypoint position.

### Data Analysis

The mean synchronization values for the right and left feet were each compared between the synchronous and asynchronous conditions using a two-sample t-test assuming unequal variances using Microsoft Excel.^43^ Statistical significance was set to 0.05.

## Results

For the synchronous condition, 23.3% and 15% of the frames for keypoint 22 and 19 respectively were removed due to low OpenPose confidence values and interpolated. For the asynchronous condition, 15.3% and 12.7% of the frames for keypoint 22 and 19 respectively were removed and interpolated.

Figure 3 illustrates the detected foot strikes and downbeats for the left and right feet in the synchronous and asynchronous conditions. The mean (standard deviation) synchronization value for the right foot was significantly lower in the synchronous condition (0.25(0.12) sec) than the asynchronous condition (0.86 (0.56) sec) (*t*(28) = −5.6418, *p*<0.001). Similarly, the mean synchronization value for the left foot was significantly lower in the synchronous condition (0.13 (0.08) sec) than the asynchronous condition (0.91 (0.57) (*t*(27) = −7.0508, *p* < 0.001)

## Discussion

Using the OpenPose library alongside Python, MATLAB, and some basic audio production knowledge, we have developed a cost-efficient method for quantifying movement synchrony with music during salsa dancing. We were able to track and quantify a dancer’s movements while performing a basic salsa step. We then compared this information to the downbeats of the salsa song to which the dancer performed the movement to determine how close the timing of the salsa steps was to the downbeat of the song. Being able to quantify movement synchrony while dancing is useful for future work that will explore the relationship between the physical and psychosocial benefits of dance and the capacity for movement synchronization to music.

Potential problems that occur with our methods include background noise and low OpenPose keypoint confidence values. Due to this fact, our method uses audio waveforms to sync up the audio and video information, however, background noise can interfere with waveform analysis. We attempted to address this by positioning the speaker playing the salsa track as close as possible to the webcam recoding the dance video. Another concern is that confidence values for some keypoints can be very low depending on angles and the type of movement recorded. For example, these low confidence values can indicate a dropped frame or OpenPose confusing body parts. We attempted to address this by removing visual clutter in the frame, properly lighting the participant during trials, removing points with low confidence, and interpolating between the points with higher confidence.

OpenPose has already been used in gait analysis research.^34,44,45^ We believe that our work here adds to the research on using accessible and affordable deep-learning-based keypoint estimators in the analysis of complex movements such as dance.

## Acknowledgements

This work was supported by a Collaboratives Health Research Project (CHRP) grant jointly funded by the National Science and Engineering Research Council and the Canadian Institutes of Health Research.

